# Chromatin attachment to the nuclear matrix represses hypocotyl elongation in *Arabidopsis thaliana*

**DOI:** 10.1101/2023.07.05.547753

**Authors:** Linhao Xu, Shiwei Zheng, Katja Witzel, Eveline Van De Slijke, Alexandra Baekelandt, Evelien Mylle, Daniel Van Damme, Jinping Cheng, Geert De Jaeger, Dirk Inze, Hua Jiang

**Affiliations:** Leibniz Institute of Plant Genetics and Crop Plant Research, Gatersleben, 06466, Germany; Leibniz Institute of Vegetable and Ornamental Crops, Großbeeren, 14979, Germany; Department of Plant Biotechnology and Bioinformatics, Ghent University, Ghent 9052, Belgium; VIB Center for Plant Systems Biology, Ghent 9052, Belgium

**Keywords:** hypocotyl growth, AT-hook, the nuclear matrix, chromatin, histone acetylation

## Abstract

The nuclear matrix is a nuclear compartment that has diverse functions in chromatin regulation and transcription. However, how this structure influences epigenetic modifications and gene expression in plants is largely unknown. In this study, we showed that a nuclear matrix binding protein, AHL22, together with the two transcriptional repressors FRS7 and FRS12, regulates hypocotyl elongation by suppressing the expression of a group of genes known as *SMALL AUXIN UP RNAs* (*SAURs*) in *Arabidopsis thaliana*. The transcriptional repression of *SAURs* depends on their attachment to the nuclear matrix. The AHL22 complex not only brings these SAURs, which contained matrix attachment regions (MARs), to the nuclear matrix, but it also recruits the histone deacetylase HDA15 to the *SAUR* loci. This leads to the removal of H3 acetylation at the *SAUR* loci and the suppression of hypocotyl elongation. Taken together, our results indicate that MAR-binding proteins act as a hub for chromatin and epigenetic regulators. Moreover, we present a novel mechanism by which nuclear matrix attachment to chromatin regulates histone modifications, transcription, and hypocotyl elongation.

## Introduction

The nuclear matrix provides a supporting scaffold for the orderly compaction of genomic DNA. It was first identified through biochemical isolation of a nuclear fraction in eukaryotic cells ^1^. This structure consists of matrix attachment regions (MARs) on the chromatin and nuclear matrix binding proteins that attach to these MARs ^2^.

MARs are short DNA sequences in diverse eukaryotic nuclear genomes, including mammals and plants ^3, 4^. These DNA sequences are notable for their richness in AT sequences that likely narrow the minor DNA groove ^5, 6^. In mammals, MARs are essential for defining structural units of chromatin by binding to the nuclear matrix and organizing the chromatin into distinct loop domains ^7,8,9^, leading to positive or negative transcriptional effects ^4^. In Arabidopsis (*Arabidopsis thaliana*), a preference for transcription start sites (TSS), enrichment of poly (dA:dT) tracts, and increased expression of nearby genes are observed for MARs identified on chromosome 4 using a tilling array ^10^. However, the distribution of MARs and their association with transcriptional outcomes in the entire genome is still unclear in Arabidopsis and other plants.

MAR-binding proteins recruit MARs to the nuclear matrix ^2^. One group of MAR-binding proteins contains an AT-hook motif, which can specifically bind to the AT-rich DNA sequence typical of MARs ^11, 12^. They also interact with other proteins and RNA to form larger complexes, serving as functional domains involved in multiple cellular processes such as transcriptional regulation, chromosome packaging, and development ^2, 13^. In mammals, the AT-hook motif-containing protein SATB1 (SPECIAL AT-RICH SEQUENCE-BINDING PROTEIN 1) is one of the most studied MAR-binding proteins essential for transcriptional regulation, chromatin organization, and histone modifications ^14,15,16,17,18,19^. In Arabidopsis, a group of AHL (AT-HOOK MOTIF CONTAINING NUCLEAR-LOCALIZED) proteins has been shown to bind to nuclear matrix regions ^20,21,22^. These AHLs are involved in many aspects of plant development, including hypocotyl elongation ^23^. Higher-order mutants for various *AHL*s and the dominant-negative mutant allele of *AHL29*, *sob3-6* (*suppressor of phyB4 #3-6*), show a longer hypocotyl than wild type plants ^23, 24^, potentially by suppressing the expression of auxin-related genes, such as *YUCCA8* (*YUC8*) and the *SMALL AUXIN UP-REGULATED RNA19* (*SAUR19*) subfamily ^25^. The expression of *SAUR19* subfamily genes can also be affected by signaling from the steroid phytohormone brassinosteroid (BR), indicative for crosstalk between different signaling pathways involved in AHL-dependent hypocotyl regulation ^26^. While it is known that phytohormone signaling pathways act downstream of AHLs in hypocotyl growth, a direct connection between MAR attachment to the nuclear matrix and gene expression, and how MAR attachment might influence gene expression are still largely unknown.

In this study, we showed that AHL22 interacts with the transcriptional repressors FRS7 (FAR1-RELATED SEQUENCE 7) and FRS12 in the Arabidopsis nuclear matrix. AHL22, FRS7, and FRS12 cooperatively suppressed hypocotyl elongation by regulating, among others, *SAUR* expression levels. In line with the proposed function of MAR-binding proteins, AHL22, FRS7, and FRS12 were required for MARs attaching to the nuclear matrix, including the MARs present in or nearby multiple *SAUR* loci. While Arabidopsis MARs preferentially mapped to highly expressed genes, we also identified several examples in lowly expressed genes, indicating that MAR-binding proteins function in both activation and repression of gene expression. Furthermore, the AHL22 complex recruited the histone deacetylase HDA15 at *SAUR* loci, leading to the silencing of *SAUR* genes and suppression of hypocotyl elongation. Taken together, our results reveal a novel mechanism by which nuclear matrix attachment of chromatin regulates histone modifications, transcription, and hypocotyl elongation.

## Results

### AHL22 interacts with FRS7 and FRS12 at the nuclear matrix

To understand how AHL22 regulates chromatin status, we identified interacting proteins using a tandem affinity purification (TAP) tagging strategy in an Arabidopsis PSB-D cell culture ^27^. Among the proteins reproducibly identified in both replicates as interactors for AHL22, we noticed multiple AHLs, including AHL27 (also named ESCAROLA [ESC]) and AHL29 (also named SOB3) (Table 1, *SI Appendix*, Table S1) ^23^. In addition, we also identified the transcriptional repressor FRS12 ^28^ (Table 1, *SI Appendix*, Table S1). FRS12, together with its paralog FRS7, regulate flowering time and hypocotyl growth by repressing the expression of their target genes ^28^, similar to the known function of other AHL proteins in transcriptional silencing ^25, 26^. Therefore, we selected FRS7 and FRS12 for further analysis. To confirm the interaction between AHL22 and FRS7/12, we performed bimolecular fluorescence complementation (BiFC) assays. We detected an interaction between AHL22 and FRS7 or FRS12, as evidenced by the strong fluorescence signals from reconstituted nuclear yellow fluorescent protein (YFP) (Fig 1*A*). Förster resonance energy transfer measured by Fluorescence Lifetime Imaging Microscopy (FRET-FLIM) experiments confirmed the interactions of AHL22-FRS7 and AHL22-FRS12 *in planta* (Fig 1*B*, *C*). Finally, in a co-immunoprecipitation (Co-IP) assay, AHL22 was found to co-purify with FRS7 and FRS12, further demonstrating the *in vivo* interaction between AHL22 and FRS7/12 (Fig 1*D*, *E*). Taken together, the above findings support the conclusion that AHL22 interacts with FRS7 and FRS12.

**Fig. 1.**
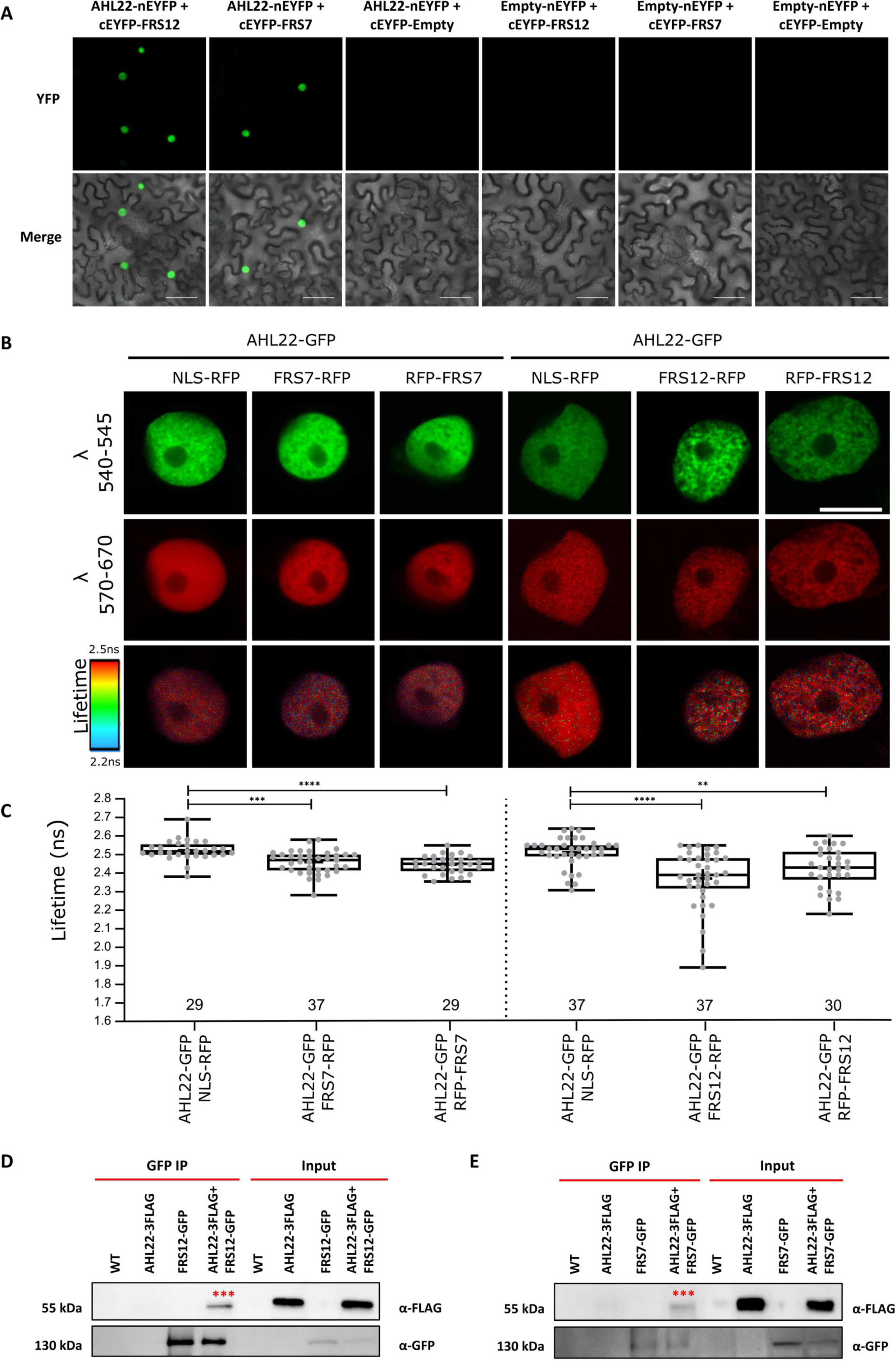
AHL22 interacts with FRS7 and FRS12. (A). Bimolecular fluorescence complementation (BiFC) analysis of AHL22 and FRS7 or FRS12 interaction *in planta*. AHL22 and FRS7 or FRS12 fused with pSITE-nEYFP-N1 and pSITE-cEYFP-C1 vectors, respectively, were cotransformed into *N. benthamiana*. Scale bar= 50 μm. (B) FRET-FLIM analysis of AHL22 and FRS7 or FRS12 interaction *in planta*. From top to bottom, representative images of nuclei, expresseing various GFP and/or RFP fused proteins in epidermal *N. benthamiana* cells, captured between wavelengths 540-545 nm and between 570-670 nm as well as the corresponding donor fluorescence lifetime calibrated to the adjacent color bar. The respective construct combinations are indicated. Scale bar = 10 um. (C) GFP donor fluorescence lifetime quantifications of the indicated construct combinations presented in a min to max box and whiskers plot showing the median (line in the middle of the box), mean (+ in the box), min and max (whiskers), individual points (gray circle) (n>=18). Statistical significance was calculated using a Welch’s Anova test combined with Dunnett’s T3 multiple comparisons test. P-values are represented as **** (p<0.0001), *** (p<0.001), ** (p<0.01). The experiment was performed in two independent infiltrations and the exact number of measurements is indicated per construct combination. (D,E). Co-immunoprecipitation assays showing interactions between AHL22 and FRS12 or FRS7. AHL22-3FLAG and FRS12-GFP or FRS7-GFP were expressed in *N. benthamiana*. WT, empty leaves. IP, immunoprecipitation. Asterisks indicate targeted proteins.

**Table 1.** Tandem affinity purification-Mass spectrometric analysis of AHL22 Expanded View Figure legends.

| Protein | Annotation | % coverage (rep 1) | No. of protein sequences (rep 1) | Protein Score (rep 1) | % coverage (rep 2) | No. of protein sequences (rep 2) | Protein Score (rep 2) |
| --- | --- | --- | --- | --- | --- | --- | --- |
| AHL22 | AT2G45430 | 12 | 3 | 241 | 21.5 | 5 | 298 |
| AHL15 | AT3G55560 | 49 | 10 | 934 | 52.3 | 11 | 1223 |
| AHL17 | AT5G49700 | 44.2 | 6 | 476 | 20.7 | 4 | 331 |
| AHL13 | AT4G17950 | 18.7 | 5 | 358 | 21.2 | 5 | 305 |
| AHL19 | AT3G04570 | 25.1 | 4 | 306 | 21 | 4 | 255 |
| SPFH | AT5G51570 | 27.7 | 5 | 231 | 19.5 | 4 | 273 |
| AHL27 | AT1G20900 | 19.6 | 3 | 221 | 19.6 | 3 | 197 |
| AHL3 | AT4G25320 | 15.3 | 4 | 215 | 16.3 | 4 | 297 |
| AHL1 | AT4G12080 | 29.2 | 4 | 199 | 39.3 | 8 | 425 |
| AHL4 | AT5G51590 | 14.8 | 3 | 121 | 6.9 | 2 | 90 |
| HTA7 | AT5G27670 | 34.7 | 2 | 119 | 42.7 | 3 | 151 |
| FRS12 | AT5G18960 | 9.1 | 3 | 98 | 4.9 | 2 | 78 |
| VRN1 | AT3G18990 | 8.8 | 3 | 93 | 8.5 | 2 | 113 |
| AHL29 | AT1G76500 | 10.6 | 2 | 88 | 20.5 | 4 | 147 |

As AHLs have been proposed to be MAR-binding proteins ^20,21,22, 29^, we hypothesized that AHL22 might interact with FRS7 and FRS12 at the nuclear matrix. To test this hypothesis, we isolated the Arabidopsis nuclear matrix fraction, in which other type of nuclear-localized proteins are removed by high concentration salf solution (Fig S1) ^30^. We identified constituent of MAR proteins by liquid chromatography-mass spectrometry (LC-MS) (Fig 2*A*, *SI Appendix*, Table S2). We thereby identified 665 nuclear matrix proteins, including the conserved nuclear matrix proteins SUN (Sad1/UNC-84) proteins AtSUN1 and AtSUN2 ^31, 32^ (Fig 2*B*). Moreover, we detected several proteins related to chromatin and transcription (*SI Appendix*, Table S2), supporting a tight association between the nuclear matrix and gene expression. Satisfyingly, we also detected multiple AHL and FRS proteins among the 665 nuclear matrix proteins (Fig 2*B*). AHL22, FRS7 and FRS12 were all among the list of proteins identified (Fig 2*B*). Hence, we conclude that FRS7 and FRS12 are likely nuclear matrix proteins that can interact with AHL22 at the nuclear matrix.

**Fig. 2.**
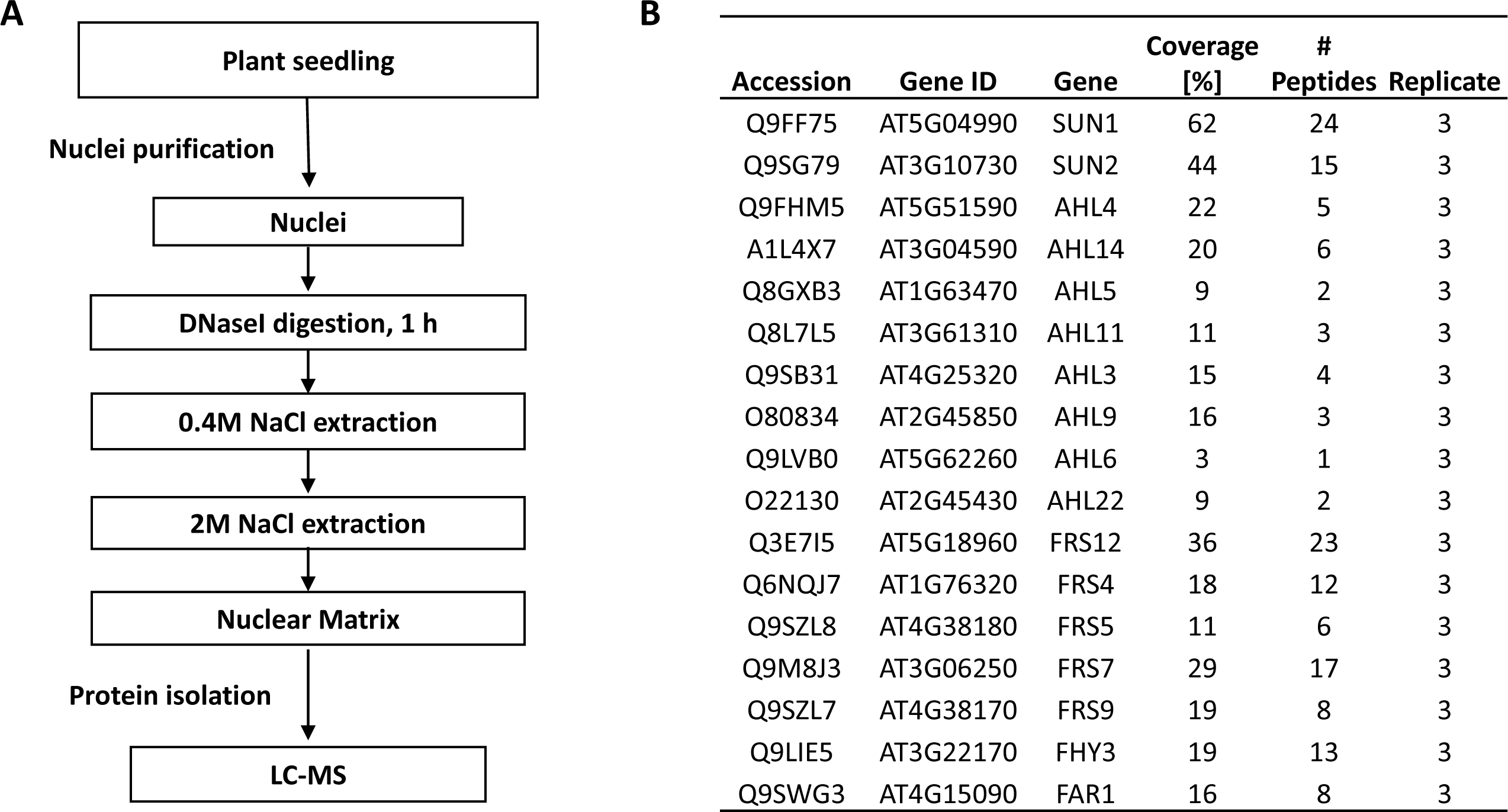
Composition of the nuclear matrix-associated proteins. (A). Flow diagram representing isolation of nuclear matrix-associated proteins (B). Identified AHL and FRS-related proteins from the LC-MS analysis, based on three biological experiments, and each experiment includes two technical replicates.

### AHL22 cooperates with FRS7 and FRS12 to repress hypocotyl elongation

FRS7, FRS12, and other AHL proteins participate in the regulation of hypocotyl elongation ^23,24,25,26, 28^, prompting us to ask if they did so cooperatively. We therefore generated an *ahl22 frs7 frs12* triple mutant. The triple mutant showed a significantly longer hypocotyl than the wild-type Col-0 (WT), the *ahl22* single mutant, or the *frs7 fr12* double mutant under LD conditions (16h 22 μmol·m−2·s−1 continuous white light, 8h dark) (*p-*value<0.01, Fig 3*A*, *B*). In agreement with previous results ^25, 26^, the dominant-negative mutant allele of *AHL29*, *sob3-6*, presented significantly elongated hypocotyl under the same conditions (*p-*value<0.01, Fig S2*A*, *B*). In contrast to the *ahl22 frs7 frs12* triple mutant, transgenic lines individually overexpressing *AHL22*, *FRS7*, or *FRS12* all exhibited significantly shorter hypocotyls than in Col-0 (*p-*value<0.01, Fig S2*C*, *D*). These results indicated that AHL22, FRS7, and FRS12 are likely to coordinately repress hypocotyl elongation.

**Fig. 3.**
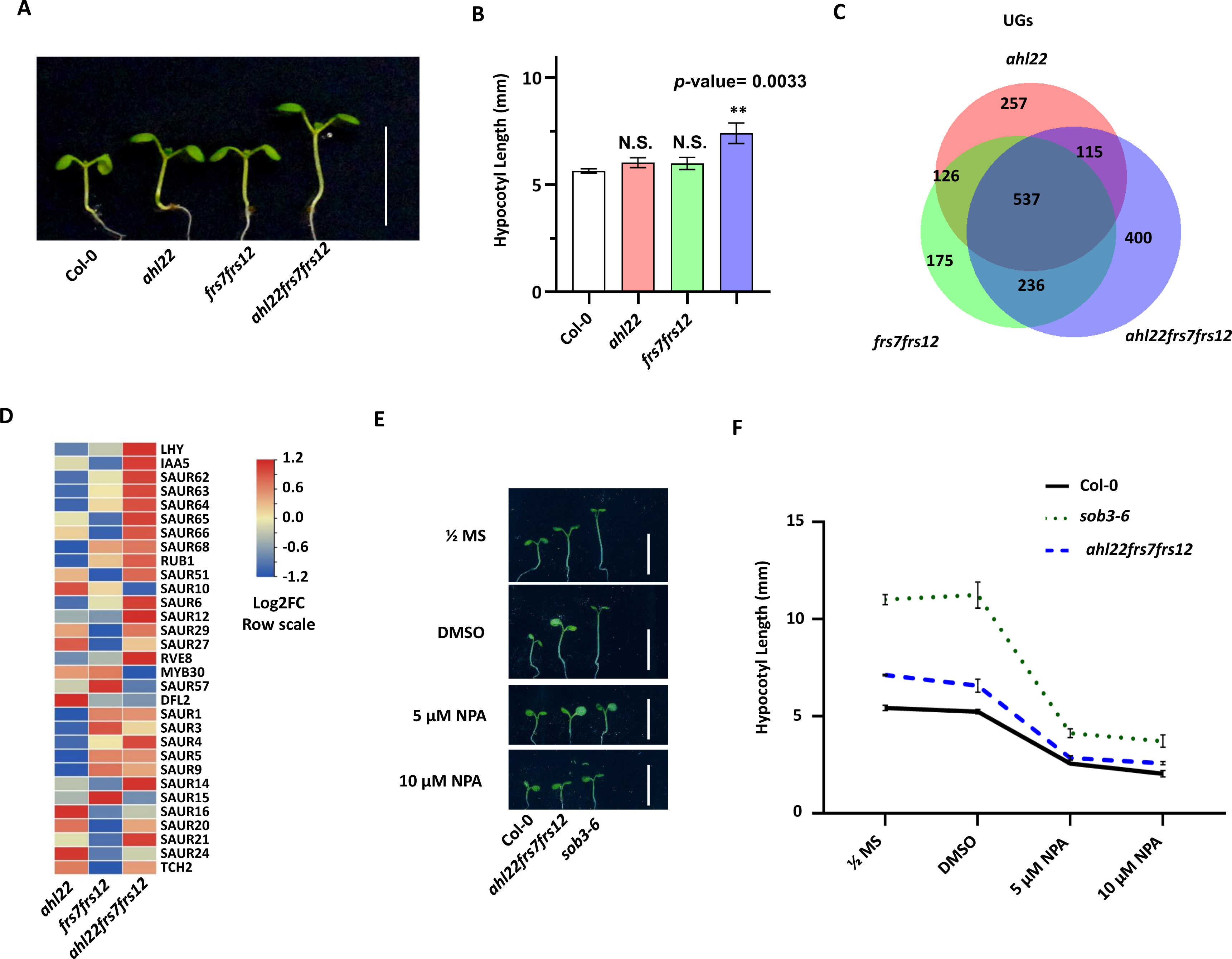
AHL22 and FRS7/FRS12 co-repress hypocotyl growth. (A). The hypocotyl phenotype of indicated lines grown vertically under LD conditions (16h at 23 °C under 22 μmol·m^−2^·s^−1^ continuous white light, 8h at 23 °C dark) for 5 days. Scale bar= 1 cm. (B). Hypocotyl lengths of indicated lines grown under LD conditions (16h at 23 °C under 22 μmol·m^−2^·s^−1^ continuous white light, 8h at 23 °C dark) for 5 days. Average length of three independent measurements ± standard deviations are shown. Each measurement with n=30 plants. (C). Venn diagrams showing overlap of upregulated genes (UGs) in *ahl22*, *frs7 frs12 and ahl22 frs7 frs12* compared with Col-0. (D). Heat map showing the relative mRNA expression levels of genes responsive to auxin in *ahl22*, *frs7 frs12 and ahl22 frs7 frs12*, Log2FC row scale was used. (E). The hypocotyl phenotype of indicated lines grown vertically under LD conditions (16h at 23 °C under 22 μmol·m^−2^·s^−1^ continuous white light, 8h at 23 °C dark) for 5 days. Scale bar= 1 cm. (F). Hypocotyl lengths of indicated lines grown under LD conditions (16h at 23 °C under 22 μmol·m^−2^·s^−1^ continuous white light, 8h at 23 °C dark) for 5 days. Average length of three independent measurements ± standard deviations are shown. Each measurement with n=30 plants. Unpaired two-tailed Student’s t-test was used to determine significance. N.S. *p value* >0.05, \**p value* ≤ 0.05, \*\**p value* ≤ 0.01, \*\*\**p value* ≤ = 0.001, \*\*\*\**p value* ≤ = 0.0001.

Given that AHL22, FRS7, and FRS12 are all transcriptional regulators ^25, 28^, we explored how they might control hypocotyl elongation via regulating gene expression. To this end, we generated transcriptome profiles for Col-0, *ahl22*, *frs7 frs12*, and *ahl22 frs7 frs12* seedlings. We identified 3155, 3108, and 3517 differentially expressed genes (DEGs, Log_2_FC > 1 or Log_2_FC < –1, with a false discovery rate [FDR] < 0.05) in *ahl22*, *frs7 frs12*, and *ahl22 frs7 frs12*, respectively, compared with Col-0 (Table S3). We detected substantial overlaps between the DEGs of *ahl22*, *frs7 frs12*, and *ahl22 frs7 frs1*2 plants (Fig 3*C*, and Fig S3*A*). Notably, UGs and DGs in *ahl22*, *frs7 frs12*, and *ahl22 frs7 frs12* were enriched in similar pathways, such as “response to auxin” and “cytoplasmic translation” for UGs, and “hydrogen peroxide catabolic process” and “response to oxidative stress” for DGs (Fig *S3B*, *C*). Among the DEGs, genes responsive to auxin, such as *SAURs*, showed higher expression levels in *ahl22 frs7 frs12* than in *ahl22* or *frs7 frs12* seedlings (Fig 3*D*), consistent with the notion that AHL27 and AHL29 suppress hypocotyl elongation by repressing genes related to auxin signaling pathways ^25, 26, 33^. In line with higher expression of *SAURs* in *ahl22 frs7 frs12*, we found that inhibiting auxin transportation by Naphthylphthalamic acid (NPA) could substantially suppress the long hypocotyl phenotype in *ahl22 frs7 frs12* (Fig 3*E*, *F*), suggesting that AHL22 and FRS7/12 indeed suppress hypocotyl elongation by repressing the auxin pathway.

To further determine the direct function of AHL and FRS proteins in gene expression, especially auxin-related genes, we explored the binding site of the above proteins. We were failed to identify peaks from the ChIP-seq experiment of AHL22-GFP. However, we found an available ChIP-seq dataset of AHL29/SOB3 in a previous publication ^34^. AHL29/SOB3 is co-precipitated with AHL22 in the IP-MS experiment (Table 1) and together regulate hypoctyl elongation with AHL22 ^23^. Therefore, we used the published dataset of AHL29/SOB3 ChIP-seq ^34^ as the representative binding site of the AHL proteins in hypocotyl regulation, along with the published dataset of FRS12^28^. Consistent with the function of SOB3 and FRS12 as transcription factors, their genome-wide distribution shared a similar pattern, with a preference for promoter-TSS regions (around 50% of all peaks) (Fig S4*A*). We compared 7,427 SOB3-bound genes ^34^ to 4,624 FRS12-bound genes ^28^, of which about 2,000 genes were shared (*p-*value=6.4e^−294^; Fig S4*B*, Table S4). These ∼2,000 genes were enriched in phytohormone pathways, including “response to auxin” and “auxin-activated signaling pathway” (Fig S4*C*), supporting the idea that AHLs and FRS7/12 proteins function as a complex to directly regulate a similar set of downstream target genes in hypocotyl elongation.

### MARs are preferentially associated with active epigenetic marks and highly expressed genes

As AHL22, FRS7, and FRS12 are likely nuclear matrix-associated proteins, we hypothesized that they might regulate transcription by recruiting chromatin to the nuclear matrix. To test this hypothesis, we first identified all genomic regions attached to the nuclear matrix by extracting the nuclear matrix and any bound DNA, followed by sequencing, called MAR-seq that was used in mammalian research, including DNA digestion with DNaseI and removal of other proteins by high salt solution (Fig S5*A*) ^35, 36^. To exclude the possibility of DNaseI digestion resulting in false positive MAR peaks, we initially compared the peaks obtained from DNaseI-digested DNA with the peaks observed after washing the digested DNA with a high salt solution (Fig S5A). We discovered that the occurrence of MAR peaks distinctly differs from the peaks observed in DNaseI-digested chromatin (Fig 4*A*). Therefore, these results confirmed that the peaks in MAR-seq is not due to the preference of DNAseI at chromatin. We identified 6754 MAR peaks in Col-0 from two replicates (Table S5, Fig S5*B*), overrepresented at two locations over protein-coding gene bodies: downstream of the TSS, and at the transcription end site (TES) (Fig 4*B*). Differ from protein-coding genes, no obvious peaks were observed over TEs (Fig 4*B*). Together, these results indicate that MARs are preferentially distributed at protein-coding genes.

**Fig. 4.**
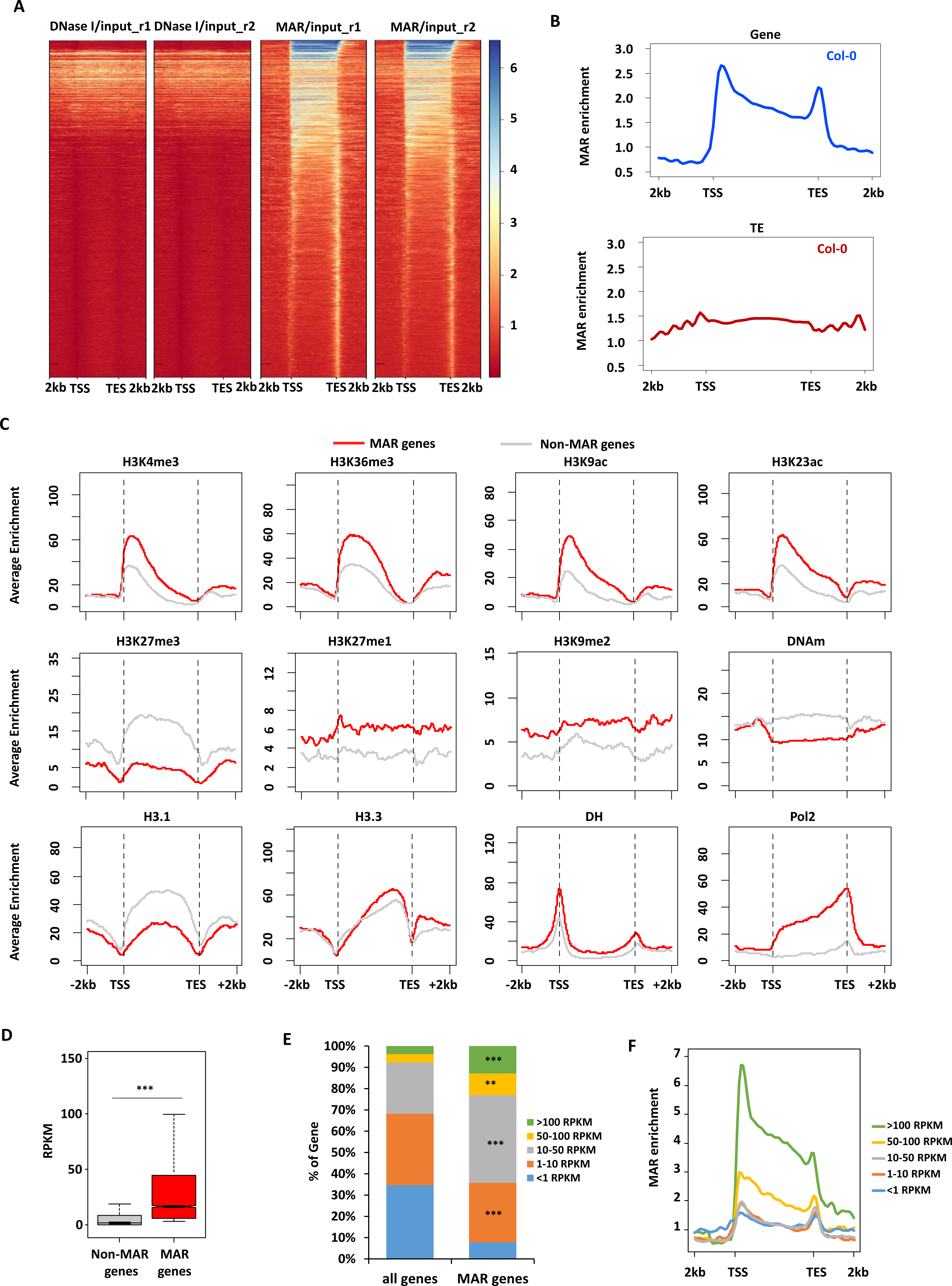
Characterization of MARs in Col-0. (A). Heatmap showing the enrichment of DNaseI/input and MAR/input peaks in genic regions. (B). Metagene plots showing the MAR enrichment of Col-0 over genes (upper) and TEs (down). Genes are scaled to align their TSSs and TTSs. Average enrichment means the percentage of regions (calculated from 100-bp windows) enriched for the respective epigenetic mark. TSS: transcription start site, TES: transcription end site. (C). Distribution of average epigenetic features across MARs over genes. (D). Box plot showing the expression level (RPKM) of non-MAR targeted and MAR targeted genes N.S. *p value* >0.05, \**p value* ≤0.05, \*\**p value* ≤ 0.01, \*\*\**p value* ≤0.001. (Kolmogorov-Smirnov test). (E). Distribution of genes over expression ranging from less than 1 RPKM, 1-10 RPKM, 10-50 RPKM, 50-100 RPKM and larger than 100 RPKM in MAR targeted genes and all genes. Fisher‘s exact test, \*\**P*<0.01, \*\*\**P*<0.001. (F). Metagene plot showing the average distribution of MAR enrichment over protein-coding genes grouped by their expression levels (RPKM).

We next determined if specific epigenetic marks were enriched in MAR-enriched genes by investigating the distribution of different epigenetic marks using published profiles ^37^. In general, MAR-enriched genes were highly correlated with active histone marks, including trimethylation at lysine 4 of histone H3 (H3K4me3), H3K36me3, and histone acetylation, with a preference for the 5’ end of protein-coding genes (Fig 4*C*). Accordingly, MAR-enriched genes were characterized with low levels of silencing marks, such as H3K27me1/3, H3K9me2, and DNA methylation (Fig 4*C*). In agreement with the presence of active histone marks, MAR-enriched genes showed significantly higher expression levels than other genes (*p-*value<0.001) (Fig 4*D*). A closer look at transcript levels indicated that the increased expression levels are driven by more highly transcriptionally active genes (with reads per kilobase of exon per million reads mapped [RPKM]>10) compared to the genome-wide pattern (Fig 4*E*). Moreover, highly transcriptionally active genes were more enriched for MARs, especially for genes with RPKM values above 50, relative to the genome-wide average (Fig 4*E*). In addition to transcriptionally active genes, some MAR-enriched genes also could be low-expressed genes (RPKM<1) (Fig 4*F*), suggesting that MAR attachment have dual functions in transcriptional regulation while it is mainly associated with gene expression.

### The AHL22-FRS7-FRS12 complex is required for MARs attaching at the nuclear matrix

To explore the role of AHL22, FRS7, and FRS12 in MAR attachment, we compared the genome-wide MAR attachment profiles in the *ahl22 frs7 frs12* triple mutant (hereafter referred to as triple mutant) to Col-0. Afterwards, we identified 5,576 genes with differential MAR peaks (FDR <0.05), of which 5,457 genes showed a lower MAR enrichment in the triple mutant (Fig 5*A*). These results indicated that the AHL22-FRS7/12 complex is required for MAR binding to the nuclear matrix. In contrast to genes with differential MAR peaks, many genes showed similar MAR attachment in WT and mutants (examples in Fig S*6*), suggesting the functional redundancy among different MAR-associated proteins.

**Fig. 5.**
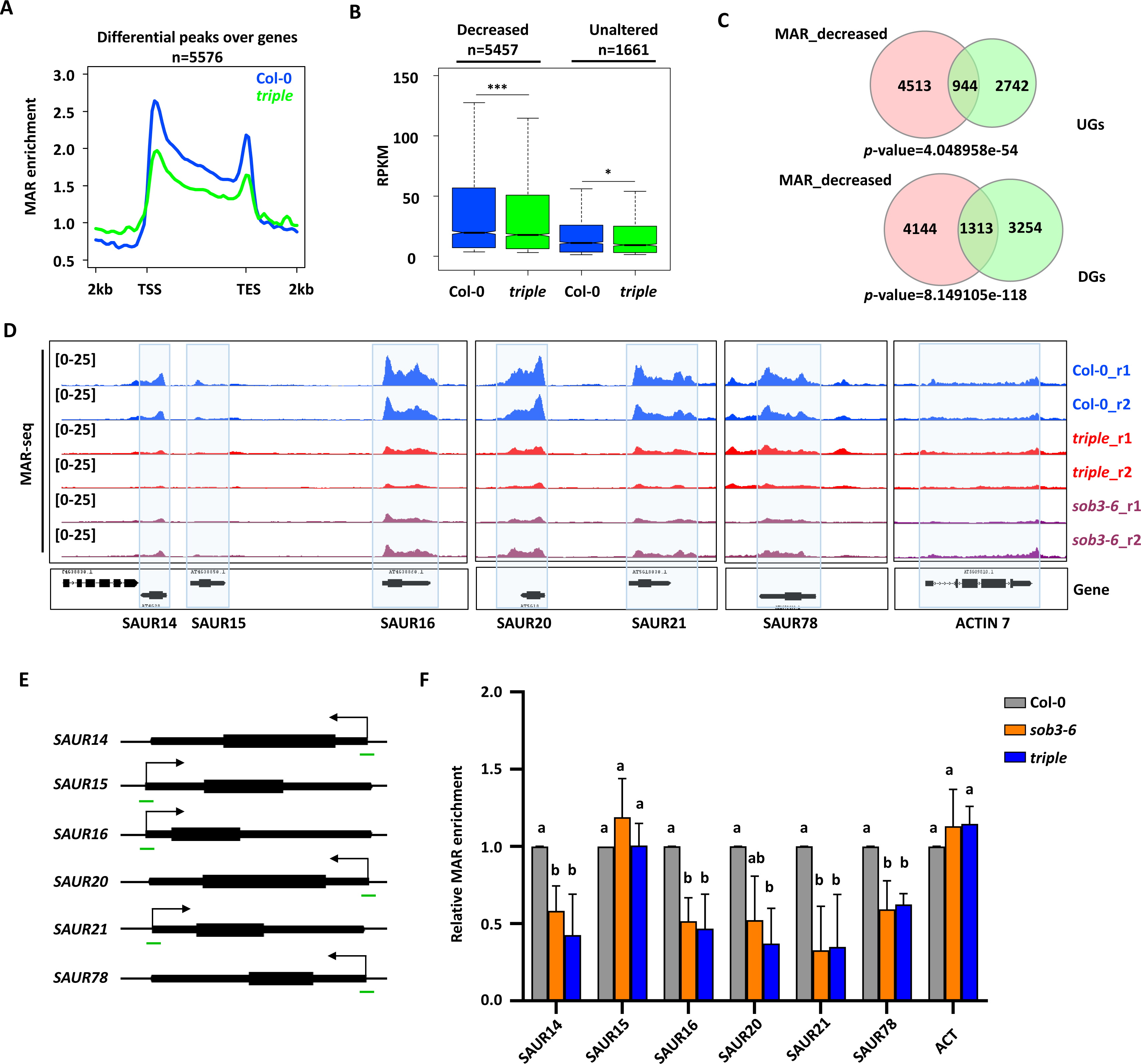
The AHL22-FRS7-FRS12 complex is required for MAR attachment. (A). Metagene plot showing the MAR enrichment of all differential peaks (n=5,576) in the *triple* mutant compared with Col-0. TSS: transcription start site, TES: transcription end site. (B). Box plot showing the expression level (RPKM) of MAR decreased (n=5,457) and MAR unaltered (n=1,661) genes. N.S. *p value* >0.05, \**p value* ≤0.05, \*\**p value* ≤ 0.01, \*\*\**p value* ≤0.001. (Kolmogorov-Smirnov test). (C). Venn diagrams showing overlap of MAR decreased genes and UGs (A) or DGs (B) identified in the *triple* mutant compared with Col-0 (DEGs, Log2FC > 0 or Log2FC < 0, with a false discovery rate [FDR] < 0.05). Significance was tested using a hypergeometric test. (D). Genome browser views of two replicates of auxin response genes indicated in blue box in Col-0, *triple* mutant and *sob3-6*. ACTIN 7 was used as control. r1/r2 (replicate1/2). (E). Schematic structures of selected genes. Arrows indicate transcription start sites (TSS). Green lines indicate regions examined by MAR-qPCR. (F). Relative MAR enrichment at selected genes determined by MAR-qPCR in Col-0, *sob3-6* and *triple*. Values are means ± SD of three biological repeats. The significance of differences at each gene was tested using one-way ANOVA with Tukey’s test (*P* < 0.05), and different letters indicate statistically significant differences.

To investigate the distribution of these decreased MAR peaks in the triple mutant as a function of gene length, we divided the 5,457 genes with a lower MAR enrichment in the triple mutant into six groups based on their size and plotted the corresponding MAR distribution over the gene body. Importantly, the distribution of decreased MAR peaks was comparable across all gene sizes (Fig S7*A*), indicating that the loss of MARs is global rather than local. However, genes shorter than 2 kb in length were highly enriched in MARs compared to those longer than 2 kb and largely contributed to the overall decrease of MAR enrichment in the triple mutant (Fig 5*B,* Fig S7*B*). Thus, small genes appear to have more MARs and contribute to the overall loss of MAR enrichment in the triple mutant.

Given that MAR-enriched genes are mainly associated with active epigenetic marks and highly transcribed genes (Fig 4*C-E*), we hypothesized that lowering MAR attachment in *ahl22 frs7 frs12* at the targeted genes might cause their transcriptional repression. Indeed, genes with fewer MARs had significantly lower expression levels in the triple mutant compared to Col-0 (Fig 5*B,* Kolmogorov-Smirnov test, *P*<0.001). However, when comparing genes with decreased peaks with the down-regulated genes (DGs) and up-regulated genes (UGs), we found both MAR-decreased genes were significantly overlapped with both DGs and UGs (Fig 5*C*). Hence, AHL22-FRS7/12-mediated MAR attaxchment is associated with both gene expression and repression.

Arabidopsis AHL27 and AHL29 inhibit hypocotyl elongation by repressing the expression of genes involved in both auxin and BR signaling pathways, including *YUC8*, *YUC9*, and members of the *SAUR19* subfamily ^25, 26, 33^. Similarly, upregulated genes in the triple mutant with lower MAR peaks were significantly enriched in genes responsive to auxin (Fig S8). In our analyses, several *SAUR* genes showed a decreased MAR enrichment and a higher expression level in the triple mutant compared to Col-0, including *SAUR14/15/16*, *SAUR20/21*, and *SAUR78* (Fig. 5*E*). We also observed a decline in MAR enrichment in the *sob3-6* mutant (Fig. 5*D*), further supporting the idea that AHL proteins are required for MAR attachment to the nuclear matrix. Moreover, these *SAUR* genes are short genes of less than 1 kb in length, which were highly enriched in MARs compared to longer genes and exhibited a noticeable decrease of MAR occupancy in the triple mutant (Fig S7). We validated these results by RT-qPCR at selected *SAUR* genes: relative to Col-0, both the triple mutant and *sob3-6* displayed significantly lower occupancy of MARs over *SAUR* genes, except for *SAUR15*, which might be due to the low overall MAR enrichment over this locus (Fig. 5*E, F*). Taken together, these data indicate that the AHL22-FRS7-FRS12 complex is responsible for MAR attachment at auxin-response genes, suppressing the expression of multiple *SAUR* genes to inhibit hypocotyl elongation.

### AHL22 recruits HDA15 to repress auxin-responsive genes

We further explored how MARs attaching to the nuclear matrix might influence transcription. Multiple AHLs are known to interact with histone deacetylases, which are essential for transcriptional silencing ^29^. Hence, we selected HDA15, whose loss of function also results in longer hypocotyls ^38^, as a candidate interactor of AHL22. The *hda15-1* mutant displayed a significantly longer hypocotyl phenotype in the same growth conditions as the triple and *sob3-6* mutants (Fig 6*A*, *B*), supporting a role for AHL22 and HDA15 in co-regulating gene expression to shape hypocotyl elongation. In line with the phenotype of longer hypocotyl, BiFC assays revealed a strong interaction between AHL22 and HDA15, with around 40% of the nuclei showing fluorescence from YFP (Fig. 6*C*, *D*). Moreover, FRET-FLIM confirmed the interaction between AHL22 and HDA15 *in planta* (Fig. 6*E*, *F*). To further test this idea, we compared the transcriptome profiles of the triple mutant and that of *hda15-1*. We identified 1,179 DEGs (Log_2_FC > 1 or Log_2_FC < –1, FDR < 0.05) in *hda15-1* compared with Col-0, with 556 DGs and 623 UGs (Table S3). These DGs and UGs showed a significant overlap with the DGs and UGs in the triple mutant, respectively (Fig 7*A*), indicating that AHL22-FRS7-FRS12 and HDA15 are likely to act together in transcriptional regulation. Moreover, manual inspection of RNA-seq datasets from the *hda15-1* and *ahl22 frs7 frs12* mutants showed that HDA15 and the AHL22-FRS7-FRS12 complex co-repress a number of genes involved in auxin responses, such as *SAUR* family genes. RT-qPCR analysis confirmed that the expression of *SAUR6*, *SAUR14*, *SAUR15*, *SAUR16*, *SAUR20*, *SAUR21*, *SAUR51*, and *SAUR78* is significantly higher in *hda15-1* compared with the wild type (Fig 7*B*), as in the triple mutant, supporting the notion that HDA15 and the AHL22-FRS7-FRS12 complex exert a repressive function for hypocotyl elongation via the auxin signaling pathway.

**Fig. 6.**
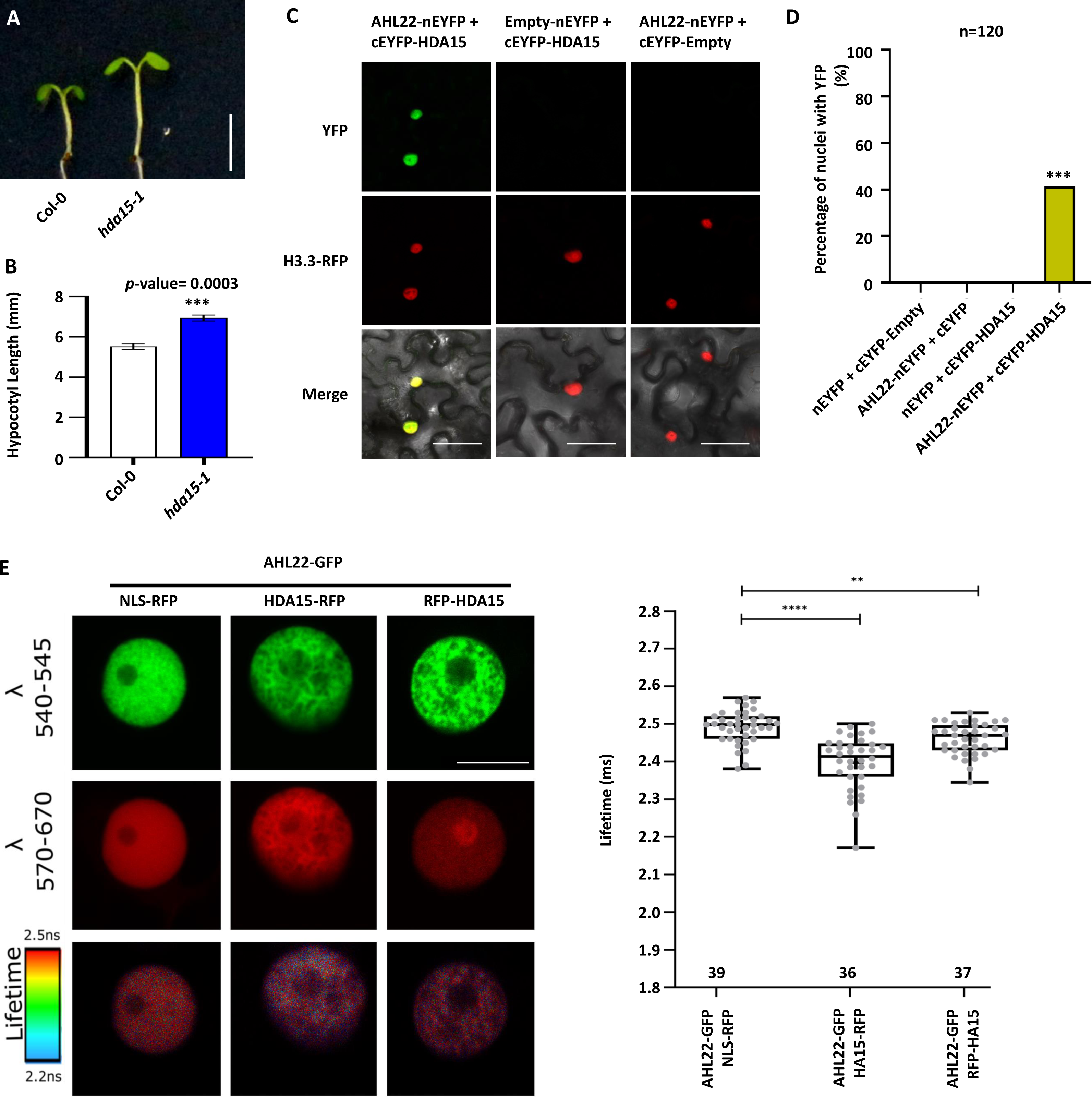
AHL22 cooperates with HDA15 to repress the expression of auxin responsive genes. (A). The hypocotyl phenotype of *hda15-1* grown vertically under LD conditions (16h at 23 °C under 22 μmol·m^−2^·s^−1^ continuous white light, 8h at 23 °C dark) for 5 days. Scale bar= 0.5 cm. (B). Hypocotyl lengths of Col-0 and *hda15-1*. Average length of three independent measurements ± standard deviations are shown. Each measurement with n=30 plants. Unpaired two-tailed Student’s t-test was used. N.S. *p value* >0.05, \**p value* ≤ 0.05, \*\**p value* ≤ 0.01, \*\*\**p value* ≤ = 0.001, \*\*\*\**p value* ≤ = 0.0001. (C). Bimolecular fluorescence complementation (BiFC) analysis of AHL22 and HDA15 interaction *in vivo*. AHL22 and HDA15 fused with pSITE-nEYFP-N1 and pSITE-cEYFP-C1 vectors, respectively, were cotransformed into *N. benthamiana*. H3.3 RFP was used to indicate the nuclei. Scale bar = 50 um. (D). Quantification of the nuclei with both YFP and RFP signal from different pairwise of BiFC experiments. n=120. Fisher‘s exact test, \*\**P*<0.01, \*\*\**P*<0.001. (E) GFP donor fluorescence lifetime quantifications of indicated construct combinations are presented in a min to max box and whiskers plot showing the median (line in the middle of the box), mean (+ in the box), min and max (whiskers), individual points (gray circle) (n>=18). Significance was calculated using a Welch’s Anova test combined with Dunnett’s T3 multiple comparisons test. P-values are represented as **** (p<0.0001), *** (p<0.001), ** (p<0.01). The experiment was performed in two independent infiltrations and the exact number of measurements was indicated per construct combination.

**Fig. 7.**
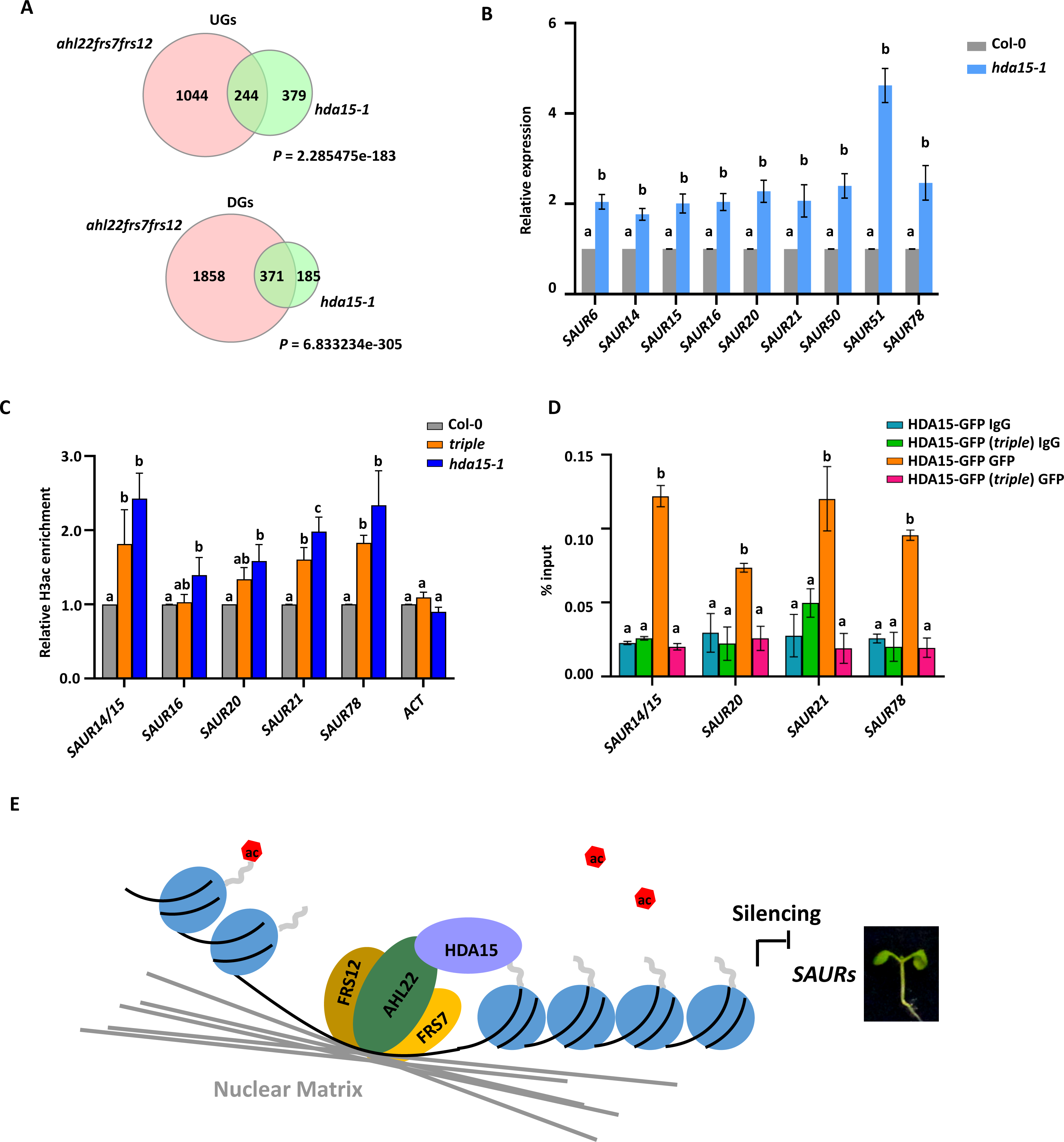
AHL22 recruits HDA15 to remove histone acetylation. (A) Venn diagrams showing overlap of UGs and DGs in *ahl22frs7frs12* and *hda15-1*. Significance was tested using a hypergeometric test. (B). Relative expression level of the selected genes by qRT-PCR from 5-d-old whole plants in *hda15-1*. Values are means ± SD of three biological repeats. The significance of differences at each gene was tested using one-way ANOVA with Tukey’s test (*P* < 0.05), and different letters indicate statistically significant differences. (C). Relative H3ac enrichment at selected genes determined by ChIP-qPCR in Col-0, *triple* and *hda15-1* plants. Values are means ± SD of three biological repeats. The significance of differences at each gene was tested using one-way ANOVA with Tukey’s test (*P* < 0.05), and different letters indicate statistically significant differences. (D). IgG and HDA15-GFP enrichment levels at selected genes in indicated lines. The amounts of immunoprecipitated DNA fragments were quantified by qPCR and normalized to input DNA. Values are means ± SD of three biological repeats. The significance of differences at each gene was tested using one-way ANOVA with Tukey’s test (*P* < 0.05), and different letters indicate statistically significant differences. (E) Proposed working model. AHL22 interacts with FRS7 and FRS12 at the nuclear matrix and recruits chromatin regions and histone deacetylases, HDA15, to the nuclear matrix and silence the target genes by reducing H3ac level to repress hypocotyl growth.

As a histone deacetylase, HDA15 negatively controls hypocotyl elongation by removing histone acetylation marks ^38^. To examine histone acetylation levels, we first performed an immunoblot on total protein extracts from Col-0, the triple mutant, and *hda15-1*, and observed a slight increase of H3 acetylation (H3ac) in the *hda15-1* mutant but not in the triple mutant (Fig S9). We speculate that the AHL22-FRS7-FRS12 complex only influences H3 acetylation marks at MARs, but not across the entire genome, as HDA15 might. We then assessed the H3ac levels by chromatin immunoprecipitation followed by quantitative PCR (ChIP-qPCR) over the promoter regions of selected *SAUR* genes, *SAUR14/15*, *SAUR20*, *SAUR21*, and *SAUR78*. We detected a significant increase of H3ac levels at the promoter regions of these genes in both *hda15-1* and the triple mutant compared to Col-0 (Fig 7*C*). In line with increased H3 acetylation levels, the binding of HDA15-GFP at the above SAUR loci was decreased in the triple mutant compared to WT (Fig 7*D*). Together, these results support that the AHL22-FRS7-FRS12 complex represses the expression of auxin-response genes by recruiting HDA15 and removing H3 acetylation at targeted loci in hypocotyl growth.

## Discussion

The data presented here demonstrate that AHL22, together with FRS7 and FRS12, regulates gene expression modulating hypocotyl elongation in light-grown Arabidopsis seedlings. Multiple AHLs repress hypocotyl growth, such as AHL27 and AHL29. Here, we show that AHL22 interacts with AHL27 and AHL29, suggesting that AHLs usually function as heteromers, in agreement with previous results described for AHL29 ^23^ and AHL10 ^39^. Moreover, we identified the non-AHL proteins FRS7 and FRS12 as interacting partners of AHL22. The *ahl22 frs7 frs12* triple mutant had a longer hypocotyl relative to the wild type, as also observed in higher-order *ahl* mutants. By contrast, individually overexpressing *AHL22*, *FRS7*, or *FRS12* shortened the hypocotyl compared to the wild type, suggesting that AHL22, FRS7, and FRS12 together regulate hypocotyl growth. We also demonstrated that similar to AHL29, the AHL22-FRS7-FRS12 complex regulates hypocotyl elongation by suppressing the expression of a group of *SAUR*s in the auxin signaling pathway. Hence, in addition to multiple AHLs, our results show that another type of MAR-binding protein acts together with AHLs at the nuclear matrix to inhibit hypocotyl growth, namely FRS7 and FRS12.

We characterized the nuclear matrix proteome, which allowed the identification of the composition of the Arabidopsis nuclear matrix. We identified multiple AHL proteins, supporting the notion that AHL proteins act as MAR-binding proteins in the nuclear matrix. We also identified multiple FRS proteins, including FRS7 and FRS12. The occurrence of other FRSs in the nuclear matrix indicated that FRS family members may commonly function as MAR binding proteins. In addition to AHL, FRS, and other chromatin- or transcription-related proteins, we also noticed multiple RNA binding proteins among the identified nuclear matrix proteins (Fig S11, Table S2), suggesting that the nuclear matrix is also a place of RNA regulation. There is at least one known example of this in mammals. The nuclear matrix protein Matrin3 can bind to RNA transcripts and regulate alternative splicing in Hela cells, besides its role as a MAR-DNA binding protein ^40^. Consistent with this example, a type of extended AT-hook motif was proposed to bind to RNAs ^41^. Therefore, multiple lines of evidence highlight the role of the nuclear matrix in RNA regulation in addition to binding to DNA.

Nuclear matrix proteins are associated with transcriptional silencing, such as CHAP (CDP-2/HDA-1-associated protein) from *Neurospora crassa* ^42^ and AHL10 from Arabidopsis ^39^. While H3K9me2 levels also increased at MAR regions of *FLOWERING LOCUS T* (*FT*) in *AHL22* overexpression lines ^29^, the *suvh4 suvh5 suvh6* (*su(var)3-9 homolog*) triple mutant did not show any hypocotyl phenotype (Fig S12), suggesting that the function of AHL22 in flowering time determination is independent from H3K9me2-mediated silencing in the context of hypocotyl regulation. We showed that AHL22 interacted with a histone deacetylase, HDA15, whose loss of function resulted in elongated hypocotyl phenotypes, similar as observed for the *ahl22 frs7 frs12* mutant under similar growth conditions. Moreover, we found that AHL22 physically interacted with and recruited HDA15 to the *SAUR* loci to remove H3 acetylation and silence the target genes. Therefore, our results demonstrate that the AHL22-FRS7-FRS12 complex act as a regulatory hub, recruiting both DNA and transcriptional repressors to the nuclear matrix and silence the targeted regions.

While the function of AHL22 in hypocotyl growth depends on transcriptional repression, we determined that MARs are preferentially associated with protein-coding genes but not TEs in the Arabidopsis genome. However, we detected MARs in highly expressed but also lowly expressed genes, a pattern that is similar to the distribution of MARs in other species ^43,44,45,46^. Therefore, MARs also appear to be associated with gene expression levels. Additional MAR regions in promoters have been shown to enhance the expression of transgenes, also supporting a role for the nuclear matrix in gene expression ^47, 48^. Apart from MAR regions, the MAR-binding protein AHL29 acts as both a positive and a negative regulator of transcription to regulate petiole growth ^34^, suggesting that AHLs also act as transcriptional activators. Hence, it will be interesting to determine whether and how nuclear matrix proteins or AHL proteins positively regulate gene expression during plant development.

## Materials and methods

### Plant materials and growth conditions

The *Arabidopsis thaliana* Columbia (Col-0) ecotype was used as the wild type. The following T-DNA mutants were used: *ahl22-1* (SALK_018866) and *hda15-1* (SALK_004027) were ordered from Nottingham Arabidopsis Stock Center (NASC). *sob3-6* mutant was kindly provided by Dr. Michael M. Neff and described in ^23^. The *frs7-1 frs12-1* double mutant, and overexpression lines of 35S::FRS7-HA and 35S::FRS12-HA were kindly provided by Prof. Dr. Alain Goossens and described in ^28^. Seeds were surface sterilized in a solution containing 75% (v/v) ethanol with 0.1% (v/v) Triton X-100 for 10 minutes and rinsed with 95% (v/v) ethanol three times before being sown on ½ MS plates (half-strength Murashige and Skoog medium, 1% sucrose, 1% plant agar). The plates were kept at 4°C in the dark for 5 days to synchronize germination before being moved into the chamber and grown vertically under LD conditions (16h at 23 °C under 22 μmol·m^−2^·s^−1^ continuous white light, 8h at 23 °C dark with 70% humidity) for 5 days.

### Plasmid construction and generation of transgenic plants

Full-length CDS with or without stop codon of *AHL22* (AT2G45430), *HDA15* (AT3G18520), *FRS7* (AT3G06250) and *FRS12* (AT5G18960) were cloned into TSK108 or pDonar201 or pENTR/D vector (Invitrogen) entry vector. For overexpression of AHL22 lines (35S::AHL22), constructs were recombined into a pB7WG2 binary vector using the LR reaction kit (Invitrogen, 11791020). The pB7WG2 destination vectors were transformed into *Agrobacterium* strain *GV3101* and then transformed into Arabidopsis Col-0 plants using the floral dip method ^49^. Transgenic plants were selected on ½ MS plates containing 30 μg·L^−1^ Basta. All primers used in this study are listed in Table S6.

### Hypocotyl Measurement

After moving the plates into the growth chamber, plated seeds were checked after 2 days, and only those with synchronized germination were included for the phenotypic analysis. 5-day-old seedlings were then scanned, and the hypocotyl lengths were quantified using the ImageJ software. Three independent measurements were taken with n=30 in each measurement. Statistical analyses were performed by unpaired two-tailed Student’s t-test.

### Tandem Affinity Purification (TAP)

Cloning of AHL22 N-terminal GS^rhino^ tag ^27^ fusion under control of the constitutive cauliflower tobacco mosaic virus 35S promoter and transformation of Arabidopsis cell suspension culture (PSB-D) with direct selection in liquid medium was carried out as previously described ^50^. TAP experiments were performed in duplo with 200 mg of total protein extract as input as described in Van Leene et al., 2015. Bound proteins were digested on-bead after a final wash with 500 µL 50 mM NH_4_HCO_3_ (pH 8.0). Beads were incubated with 1 µg Trypsin/Lys-C in 50 µL 50 mM NH_4_OH and incubated at 37°C for 4 h in a thermomixer at 800 rpm. Next, the digest was separated from the beads, an extra 0.5 µg Trypsin/Lys-C was added and the digest was further incubated overnight at 37°C. Finally, the digest was centrifuged at 20800 rcf in an Eppendorf centrifuge for 5 min, the supernatant was transferred to a new 1.5 mL Eppendorf tube, and the peptides were dried in a Speedvac and stored at −20°C until MS analysis. Co-purified proteins were identified by mass spectrometry using a Q Exactive mass spectrometer (ThermoFisher Scientific) using standard procedures ^51^. Proteins with at least two matched high confident peptides in at least 2 experiments in the dataset were retained. Background proteins were filtered out based on frequency of occurrence of the co-purified proteins in a large dataset containing 543 TAP experiments using 115 different baits ^27^. True interactors that might have been filtered out because of their presence in the list of non-specific proteins were retained by means of semi-quantitative analysis using the average normalized spectral abundance factors (NSAF) of the identified proteins ^27^.

### Bimolecular fluorescence complementation (BiFC)

The full-length coding sequences of AHL22, FRS7, FRS12 and HDA15 were cloned into the pSITE-nEYFP-N1 or pSITE-cEYFP-C1 vectors with the 35S promoter ^52^ using the LR reaction kit (Invitrogen, 11791020). Constructs were transiently expressed by *A. tumefaciens*-mediated transformation to 1-month-old *N. benthamiana* plants using an infiltration buffer composed of 10 mM MgCl_2_, 10 mM MES and 100 mM acetosyringone with OD=0.8, and the addition of a Hcpro-expressing *Agrobacterium strain* to boost protein expression. The YFP signal was observed 2-days after infiltration using a Zeiss confocal laser scanning microscope (LSM780). H3.3-RFP was used as the marker protein of the nuclei and used for quatitative analysis. We first chose the setting that all negative controls showed no YFP signal, excluding the possibility of false positive result. And then, we quatified the number of cells showing both YFP and RFP for all conbinations. The percentage indicates the strength of protein interactions. The experiment was performed in two independent infiltrations.

### Forster resonance energy transfer measured by Fluorescence Lifetime Imaging Microscopy (FRET-FLIM)

The donor fluorescence life time (FLIM) was determined by time-correlated single-photon counting (TCSPC) in *N. benthamiana* (tobacco) epidermal cells transiently expressing the proteins of interest fused to either eGFP (donor) or mRFP (acceptor). Images were collected using an LSM Upgrade Kit (PicoQuant) attached to a Fluoview FV-1000 (Olympus) confocal microscope equipped with a Super Apochromat 60x UPLSAPO water immersion objective (NA 1.2) and a TCSPC module (Timeharp 200; PicoQuant). A pulsed picosecond diode laser (PDL 800-B; PicoQuant) with an output wavelength of 440 nm at a repetition rate of 40 MHz was used for donor fluorescence excitation. A dichroic mirror DM 458/515 and a band pass filter BD520/32-25 were used to detect the emitted photons using a Single Photon Avalanche Photodiode (SPAD; PicoQuant). Laser power was adjusted to avoid average photon counting rates exceeding 10000 photons/s to prevent pulse pile up. Image acquisition was done at zoom 10 with an image size of 512 x 512 pixels corresponding to 0.041 um/pixel. A dwell time of 8 us/pixel was used for image acquisition. Samples were scanned continuously (maximum for 1 min) to obtain appropriate photon numbers (around 10000 peak counts) for reliable statistics for the fluorescence decay. Fluorescence life times were calculated using the SymPhoTime software package (v5.2.4.0; PicoQuant). Selected areas of the images corresponding to single nuclei (n ≥18 cells) were fitted by a bi-exponential reconvolution fitting. The calculated IRF (Instrument response function) was included, the background and shift IRF were fixed, and the amplitudes of the life times were set to be positive. The average instensity lifetimes τ for a series of measurements were subjected to the ROUT method (robust regression followed by outlier identificiation) (with Q set to 1%) to remove outliers (PMID: 16526949). The average intensity life times that were maintained after the outlier analysis are presented in a min to max box and whiskers plot showing the median (line in the middle of the box), mean (+ in the box), min and max (whiskers) and individual points (gray circles). Statistical significance was calculated using a Welch’s Anova test combined with Dunnett’s T3 multiple comparisons test in GraphPad Prism 9.0.0. The experiments were performed in two independent infiltrations.

### Co-immunoprecipitation (Co-IP)

To express tag-fused proteins in *N. benthamiana*, the entry vector TSK108-AHL22 fused with 3FLAG and TSK108-FRS7 or TSK108-FRS12 were recombined into a pB7WG2 and a pH7FWG2 binary vector with the 35S promoter using the LR reaction kit (Invitrogen, 11791020), respectively. *Agrobacterium* strains containing the constructs were co-infiltrated into *N. benthamiana* plant leaves. After 2–3 days, *N. benthamiana* leaves were collected with corresponding expression constructs as indicated. WT indicated *N. benthamiana* leaves without constructs. Co-IPs were performed as previously described in ^53^. Briefly, 1-2g tissues were ground in liquid nitrogen and resuspended in 5 ml of IP buffer (50 mM Tris HCl pH 7.5, 150 mM NaCl, 5 mM MgCl_2_, 0. 1 % NP 40, 0.5 mM DTT, 10 % glycerol, 1 mM PMSF, 1 µg/µL pepstatin, proteinase inhibitor cocktail (Roche, 11836145001)). The mixture was incubated on ice for 20 min and centrifuged at 4000g for 10 min at 4 °C. The supernatant was filtered through a double layer of Miracloth. The flow-through was incubated with 25 μl GFP-Trap Magnetic Agarose (Chromotek, gtma-20) overnight at 4 °C. After incubation, beads were collected and washed five times with IP buffer, 5 min each wash at 4 °C with rotation. Elution was performed by incubating beads in PBS and 4xSDS buffer at 95 °C for 10 min. Western blotting was performed with an anti-GFP (B-2) antibody (SANTA CRUZ sc-9996, 1:50) and anti-FLAG antibody (F1804, Sigma, 1:1000). Signals were detected with the ChemiDoc (BIO-RAD).

### RNA extraction and RT-qPCR analysis

Total RNA from 5-day-old whole plants was extracted using the MagMAX™ Plant RNA Isolation kit (Thermo Fisher #A33784) according to the manufacturer’s instructions. Second-strand cDNA was synthesized using the cDNA Synthesis kit (Thermo Fisher, K1612). RT-qPCR reactions contained Solis BioDyne-5x Hot FIREPol®EvaGreen®qPCR Supermix (ROX, Solis BioDyne, 08-36-00008), and the runs were performed in a QuantStudio 6 Flex Real-Time PCR System (Applied Biosystems). Three biological replicates were performed, using GADPH as the reference gene. The primers used are listed in Table S6.

### RNA-seq

Total RNA from 5-day-old whole plants was extracted using the RNeasy Plant Mini Kit (Qiagen, 74904). For RNA-seq, libraries were constructed with the VAHTS mRNA-seq V6 Library Prep Kit (Vazyme, NR604-01) according to the manufacturer’s instructions. The libraries were sequenced at Novogene (UK) via a Novaseq instrument in 150-bp paired-end mode. Two or three biological replicates were performed.

### Isolation of the nuclear matrix

The nuclear matrix was isolated as previously described in ^35^. Briefly, Arabidopsis 5-day-old whole plants grown under LD condition (16h at 23 °C under 22 μmol·m^−2^·s^−1^ continuous white light, 8h at 23 °C dark with 70% humidity) were collected and chopped in 1 ml LB01 buffer (2 mM Na_2_EDTA, 20 mM NaCl, 2 mM EDTA, 80 mM KCl, 0.5 mM spermine, 15 mM β-mercaptoethanol, 0.1% Triton X-100, pH 7.5). Another 9 ml LB01 buffer was used to wash the nuclei into 1-layer miracloth filter followed by a one-time filter through 30 μm CellTrics (Sysmex, Germany, 04-0042-2316). The nuclei mixture was centrifuged at 1500g for 10 min at 4 °C. The nuclei pellet was further wash with DNase I buffer (20 mM Tris pH 7.4, 20 mM KCl, 70 mM NaCl, 10 mM MgCl2, 0.125 mM spermidine 1 mM PMSF, 0.5% Triton-X 100) until it turned white. After centrifugation, the nuclei were resuspended in 200 μl DNase I buffer, and an aliquot of nuclei was stored for isolation and estimation of total genomic DNA as control. DNase I (Thermo Scientific, EN0525) was added and incubated at 4°C for 1 hr. Nuclei were collected by centrifugation at 3000g for 10 min at 4 °C. Digestion was followed by extraction with 0.4 M NaCl for 5 min twice on ice in Extraction Buffer (10 mM Hepes pH 7.5, 4 mM EDTA, 0.25 mM spermidine, 0.1 mM PMSF, 0.5% (v/v) Triton X-100). Followed by another two rounds of extraction with 2 M NaCl for 5 min on ice in Extraction Buffer. The final nuclear matrix pellet was washed twice with Wash Buffer (5 mM Tris pH 7.4, 20 mM KCl, 1 mM EDTA, 0.25 mM spermidine, 0.1 mM PMSF).

The nuclei from different steps as indicated were checked by 4′,6-diamidino-2-phenylindole (DAPI) staining and analyzed using an LSM780 confocal microscope (Zeiss) to validate DNA content in the nuclei (Fig S14). For protein validation, after chopping, the nuclei mixture was separated into several same-volume nuclei and followed the extraction procedure. Proteins were extracted from supernatant or pellet as indicated. All proteins were separated on 10% Tricine-SDS-PAGE gels ^54^ followed by Coomassie staining. The gel was visualized with the ChemiDoc (BIO-RAD) (Fig S14).

### Proteome analysis of the nuclear matrix

Proteins present in the nuclear matrix, isolated from three independent preparations of 1 g of plants as described above, were solubilized in using RapiGest SF (Waters, Germany), then reduced and digested following the method of Kaspar et al. ^55^. Desalting of peptides was done using Peptide Desalting Spin Columns (Pierce, Thermo Scientific, United States) following the manufacturer’s instructions. Peptides were resuspended in 2% acetonitrile/0.1% trifluoroacetic acid to a volume of 30 µl and 2 µL of protein digest was analyzed using nanoflow liquid chromatography on a Dionex UltiMate 3000 system (Thermo Scientific) coupled to a Q Exactive Plus mass spectrometer (Thermo Scientific) as described previously ^56^. Each sample was measured in duplicate. The raw files were processed using Proteome Discoverer 2.4 and Sequest HT engine (Thermo Scientific), searching the SwissProt *Arabidopsis thaliana* (TaxID=3702) database (downloaded July 2021). Precursor ion mass tolerance was set to 10 ppm and fragment ion mass tolerance was set to 0.02 Da. False discovery rate (FDR) target values for the decoy database search of peptides and proteins were set to 0.01 (strict level for highly confident identifications). Protein abundance quantification was done using the Top N average method, (N= 3). The result lists were firstly filtered and proteins were only kept for further investigation that fulfilled the following characteristics: identified by at least two peptides or by one peptide representing at least 10% protein coverage. Moreover, we further removed the proteins annotated as ribosome or other cytoplasmic localized proteins in GO analysis, which were considered as background from the nuclear matrix purification.

### MAR-qPCR and MAR-seq

DNA was extracted from the isolated nuclear matrix and the input sample. To remove RNA, RNase A (Thermo Scientific, EN0531) was added to a final concentration of 20 μg/ml and incubated for 30 min at 37°C. This was followed by digestion with 100 μg/ml Proteinase K (Ambion, AM2546) at 55°C for 1 hr. DNA was recovered by extraction with phenol:chloroform:isoamyl alcohol (25:24:1) and ethanol precipitation. The isolated input DNA was fragmented by using a Bioruptor® Plus sonication device (Diagenode) to obtain the desired DNA fragment size (enriched at 200 bp).

The amount of fragmented DNA was quantified by qPCR. Three biological replicates were performed. Statistical analyses were performed by using one-way ANOVA with Tukey’s test (P < 0.05), and different letters indicate statistically significant differences. The primers used are listed in Table S6.

Libraries were prepared using NEBNext® Ultra™ II DNA Library Prep Kit (NEB, E7645) according to the manufacturer’s instructions. The libraries were sequenced at Novogene (UK) via a Novaseq instrument in 150-bp paired-end mode. Two biological replicates were performed.

### Bioinformatic analysis

For RNA-seq analysis, Reads were mapped to the TAIR10 wild-type Arabidopsis genome with HISAT2 ^57^ in paired-end mode. Differentially expressed genes were analyzed via the Subread ^58^ and DESeq2 ^59^ R packages with a 0.05 false discovery rate (FDR). The volcano plot and heatmap of DEGs were generated using TBtools ^60^. GO term enrichment was determined at https://david.ncifcrf.gov/summary.jsp.

For MAR-seq analysis, quality control and adapter trimming of MAR-seq reads were performed with Fastp ^61^. Reads were aligned to the TAIR10 reference genome using Bowtie2. Mapped reads were deduplicated using MarkDuplicates (https://broadinstitute.github.io/picard). Coverage was estimated and normalized to 10 million reads. MAR-seq peaks in wild-type and mutants were called by SCIER2 ^62^. IGB genome browser was used to visualize the data and to generate screenshots. Boxplots and metaplots were generated with R. Kolmogorov-Smirnov tests were performed at https://scistatcalc.blogspot.com. GO term enrichment was determined at https://david.ncifcrf.gov/summary.jsp.

### Histone extraction and western blot

The histone was extracted by the EpiQuik Total Histone Extraction Kit (EPIGENTEK, OP-0006-100) and followed manufacturer recommendations, using 5-day-old whole plants. Extracted histones were separated on 10% Tricine-SDS-PAGE gels ^54^. Primary antibodies used included anti-H3 (Sigma/H9289), anti-H3ac (Millipore /06-599), and anti-H4ac (Millipore/06-866). Signals were detected with the ChemiDoc (BIO-RAD).

### ChIP-qPCR

Approximately 1 g of 5-day-old whole plant grown under LD condition (16h at 23 °C under 22 μmol·m^−2^·s^−1^ continuous white light, 8h at 23 °C dark with 70% humidity) was collected. The ChIP experiments were performed as described by ^63^. In brief, fresh tissue was cross-linked in 1x PBS (0.01% Triton, 1% formaldehyde) and vacuum infiltrated for 10 min on ice. The cross-linking was stopped by adding glycine to a final concentration of 125 mM. Cross-linked tissues were ground to a fine powder in liquid nitrogen and resuspended in 7 ml of Honda buffer (0.4 M Sucrose, 2.5% Ficoll, 5% Dextran T40, 25 mM Tris-HCl, pH 7.4, 10 mM MgCl2, 0.5% Triton X-100, 0.5 mM PMSF, 10 mM β-mercaptoethanol, proteinase inhibitor cocktail (Roche, 11836145001)). The mixture was incubated on ice for 20 min followed by two times filter through Miracloth and one time through 30 μm CellTrics (Sysmex, Germany, 04-0042-2316) and centrifuged at 1500g for 10 min at 4 °C. The pellet was resuspended in 150 μl Nuclei lysis buffer (50 mM Tris–HCl, pH 8.0, 10 mM EDTA, 1% SDS, 0.1 mM PMSF, 1 μM pepstatin A, Protease Inhibitor Cocktail) and sonicated by using a Bioruptor® Plus sonication device (Diagenode) to obtain the desired DNA fragment size (enriched at 500 bp). The following procedures were described by ^63^. Immunoprecipitations were performed with anti-H3 (Sigma, H9289), anti-H3ac (Millipore, 06-599), anti-GFP (Invitrogen, A11122) and IgG Isotype Control (Invitrogen, 026102) antibodies. After the de-crosslinking, DNA was purified according to the IPure kit v2 Kit manual (Diagenode, C03010015). The amount of immunoprecipitated DNA was quantified by qPCR. Three biological replicates were performed. Statistical analyses were performed by using one-way ANOVA with Tukey’s test (P < 0.05), and different letters indicate statistically significant differences. The primers used are listed in Table S6.

## Acknowledgments

We thank Dr. Michael M. Neff and Prof. Dr. Alain Goossens for kindly sharing seeds. We thank Prof. Chang Liu for the help in bioinformatics analysis. This research was supported by the intramural funding from IPK to HJ, and a grant from the DFG to HJ (JI 347/6-1).

## Data access

All sequencing data have been submitted to the NCBI Gene Expression Omnibus (GEO; https://www.ncbi.nlm.nih.gov/geo/) under accession number GSE215135. Proteome raw data has been deposited at MassIVE (https://massive.ucsd.edu/ ProteoSAFe/static/massive.jsp?redirect=auth) under the dataset ID MSV000090406.

## Author contributions

LX, KW, EVDS, GDJ, AB, DVD, EM, MS, DI, and JC, executed the experimental procedures. KW analyzed the composition of nuclear matrix associated proteins; EVDS and GDJ performed the IP-MS analysis for AHL22; AB and DI shared material, EM performed FLIM measurements. DVD, AB and EM analyzed data; LX and HJ analyzed the NGS data. LX and HJ performed the experimental design. LX and HJ wrote the manuscript. All authors discussed the results and commented on the manuscript.

---

**Fig. S1. Validation of extracted protein and DNA from nuclear matrix.**

(A). Coomassie-stained SDS-tricine gels of proteins during the extraction processes. M-Marker, S-Supernant, P-nuclei pellet.

(B). Confocal images and DAPI staining of nuclei pellet at three stages of nuclear matrix isolation. Scale bar= 50 μm.

**Fig. S2. Hypocotyl phenotype**

(A,C). The hypocotyl phenotype of indicated lines grown vertically under LD conditions (16h at 23 °C under 22 μmol·m^−2^·s^−1^ continuous white light, 8h at 23 °C dark) for 5 days. Scale bar= 1 cm.

(B,D). Hypocotyl lengths of indicated lines grown under LD conditions (16h at 23 °C under 22 μmol·m^−2^·s^−1^ continuous white light, 8h at 23 °C dark) for 5 days.

Average length of three independent measurements ± standard deviations are shown. Each measurement with n=30 plants. Unpaired two-tailed Student’s t-test was used to determine significance between wild type and mutants or overexpression lines. N.S. *p value* >0.05, \**p value* ≤ 0.05, \*\**p value* ≤ 0.01, \*\*\**p value* ≤ = 0.001, \*\*\*\**p value* ≤ = 0.0001.

**Fig. S3. Transcriptomic analysis in the mutants.**

(A) Venn diagram showing the overlap of down-regulated gene in corresponding genotypes; enriched biological processes of significantly UGs (A) and DGs (B) in *ahl22*, *frs7 frs12* and *ahl22 frs7 frs12* compared to Col-0. The X-axis represent negative log10 (p-value). Yellow stars indicate pathway response to auxin.

**Fig. S4. SOB3 (AHL29) and FRS7/12 co-target to phytohormone pathways.**

(A). Distribution of published SOB3 and FRS12 binding sites determined from ChIP-seq data. ‘‘Promoter-TSS’’ is defined as −1,000 bp to +100 bp in relation to the transcription start site. ‘‘TTS’’ is defined as −100 bp to +1,000 bp in relation to the transcription termination site.

(B). Venn diagram showing overlap of SOB3 BOUND and FRS12 BOUND sites. Significance was tested using a hypergeometric test.

(C). Enriched biological processes of common targeted genes (n=2050) of SOB3 BOUND and FRS12 BOUND. The X-axis represent negative log10 (*p-value*). Red stars indicate phytohormone pathways.

**Fig. S5. MAR-seq in WT and *ahl22 frs7 frs12*.**

(A) Flow diagram representing isolation of nuclear matrix and further DNA extraction. (B) Venn diagram showing the overlap of MAR peaks in WT between two replicates. (C) distribution of MAR peaks in genes, promoters, and TEs. (D) Genome browser views of a random selected region (Chr1: 5250000-5450000) showing MAR peaks in 2 replicates of Col-0. r1/r2 (replicate1/2).

**Fig. S6. Genome browser views of two replicates of genes with no difference of MAR enrichment in Col-0, *triple* and *sob3-6*. r1/r2 (replicate1/2).**

**Fig. S7. Distribution of decreased MAR at different sizes of genes.**

(A). Distribution of genes over gene size ranging from 2-3 kb, 3-4 kb, 4-5 kb and larger than 5 kb in MAR decreased genes and all genes.

(B). Metagene plot showing the average distribution of MAR enrichment over protein-coding genes grouped by gene size. TSS: transcription start site, TES: transcription end site.

**Fig. S8. Gene ontology analysis of MAR decreased genes overlapped with DEGs in the triple mutant.**

(A, B). Enriched biological processes of common genes identified from MAR decreased genes and UGs (A, n=232) and DGs (B, n=443) in the triple mutant. The X-axis represent negative log10 (*p-value*).

**Fig. S9. Western blot for H3ac, H4ac and H3 levels in total extracts from 5-d-old whole plants of Col-0, *ahl22 frs7 frs12* and *hda15-1*.**

**Fig. S10. Enriched molecular function of nuclear matrix proteins from 5-d-old Col-0 Fig. S11. Hypocotyl phenotype of *suvh456*.**

(A). The hypocotyl phenotype of indicated lines grown vertically under LD conditions (16h at 23 °C under 22 μmol·m^−2^·s^−1^ continuous white light, 8h at 23 °C dark) for 5 days. Scale bar= 1 cm.

(B). Hypocotyl lengths of indicated lines grown under LD conditions (16h at 23 °C under 22 μmol·m^−2^·s^−1^ continuous white light, 8h at 23 °C dark) for 5 days. Average length of three independent measurements ± standard deviations are shown. Each measurement with n=30 plants. Unpaired two-tailed Student’s t-test was used. N.S. *p value* >0.05, \**p value* ≤ 0.05, \*\**p value* ≤ 0.01, \*\*\**p value* ≤ = 0.001, \*\*\*\**p value* ≤ = 0.0001.

**Supplement Table S1. TAP-MS analysis of AHL22.**

**Supplement Table S2. LC-MS analysis of nuclear matrix proteins.**

**Supplement Table S3. Differentially expressed genes in *ahl22*, *frs7frs12*, *ahl22frs7frs12* and *hda15-1*.**

**Supplement Table S4. Common targets bound by SOB3 and FRS12. Supplement Table S5. MAR-seq of Col-0 and *ahl22frs7frs12*.**

**Supplement Table S6. List of Primers Used in This Study.**

